# The pentaglycine bridges of *Staphylococcus aureus* peptidoglycan are essential for cell integrity

**DOI:** 10.1101/479006

**Authors:** João M. Monteiro, Daniela Münch, Sérgio R. Filipe, Tanja Schneider, Hans-Georg Sahl, Mariana G. Pinho

## Abstract

Bacterial cells are surrounded by cell wall, whose main component is peptidoglycan (PG), a macromolecule that withstands the internal turgor of the cell. PG composition can vary considerably between species. The Gram-positive pathogen *Staphylococcus aureus* possesses highly crosslinked PG due to the presence of cross bridges containing five glycines, which are synthesised by the FemXAB protein family. FemX adds the first glycine of the cross bridge, while FemA and FemB add the second and the third, and the fourth and the fifth glycines, respectively. Of these, FemX was reported to be essential. To investigate the essentiality of FemAB, we constructed a conditional *S. aureus* mutant of the *femAB* operon. Depletion of *femAB* was lethal, with cells appearing as pseudomulticellular forms that eventually lyse due to extensive membrane rupture. This deleterious effect was mitigated by drastically increasing the osmolarity of the medium, indicating that pentaglycine crosslinks are required for *S. aureus* cells to withstand internal turgor. Despite the absence of canonical membrane targeting domains, FemA has been shown to localise at the membrane. To study its mechanism of localisation, we constructed mutants in key residues present in the putative transferase pocket and the α6 helix of FemA, possibly involved in tRNA binding. Mutations in the α6 helix led to a sharp decrease in protein activity *in vivo* and *in vitro* but did not impair correct membrane localisation, indicating that FemA activity is not required for localisation. Our data indicates that, contrarily to what was previously thought, *S. aureus* cells do not survive in the absence of a pentaglycine cross bridge.

## Introduction

*S. aureus* is one of the main pathogens responsible for life-threatening infections worldwide, particularly hospital- and community-acquired methicillin resistant *S. aureus* strains (HA-MRSA and CA-MRSA, respectively), which constitute a major challenge to antibiotic therapy^1,2^. Most of the widely used, and more potent antibiotics, target steps in the biosynthesis of peptidoglycan (PG), the core component of the bacterial wall. PG is a macromolecule composed of glycan chains, where each unit is constituted of N-acetylmuramic acid (MurNAc) and N-acetylglucosamine (GlcNAc) sugars, with a stem peptide attached to MurNAc. Glycan chains are connected (crosslinked) through flexible species-specific peptide bridges, creating a mesh-like structure that envelops the cell^3^. The structural features of PG confer both robustness and flexibility to the cell envelope, which are necessary to withstand high pressure derived from intracellular turgor^4^.

MRSA strains are resistant to β-lactams, which irreversibly acylate the transpeptidase domain of Penicillin Binding Proteins (PBPs), enzymes responsible for the last steps of PG biosynthesis^1^. In these strains, the major determinant of methicillin resistance is the acquired *mecA* gene, which encodes for PBP2A, an enzyme insensitive to β-lactam acylation^5^. However, high-level β-lactam resistance is in fact dependent on several additional elements, which were initially identified by transposon mutagenesis and termed *fem* (factor essential for methicillin resistance) or *aux* (auxiliary) genes^6,7^. Approximately 30 *fem/aux* determinants have been identified so far and most are housekeeping genes, involved in a variety of cellular processes and probably present in every *S. aureus* strain^8^. Three closely related factors -*fmhB* and the co-transcribed *femA* and *femB* genes, encode for the FemX, FemA and FemB proteins, respectively, peptidyltransferases which synthesise the pentaglycine bridges used to crosslink glycan chains in *S. aureus*^9,10^. During the inner membrane steps of PG synthesis (see Pinho et al.^11^ for a review), the Fem proteins sequentially transfer five glycine residues to the PG precursor lipid II using glycyl-charged tRNA molecules^12^. Importantly, *in vivo* and *in vitro* studies have shown that each Fem protein has strict substrate specificity: FemX adds the first glycine, FemA adds the second and the third and FemB adds the fourth and fifth glycines, and each Fem cannot substitute for another^13,14^. Although *fmhB* was shown to be an essential gene^15^, mutants carrying transposon inactivated *femA or femB* grew poorly but were viable, suggesting that *S. aureus* can survive with a PG composed of monoglycine crossbridges^9,16^. However, HPLC analysis of the PG composition in these mutants revealed an overall reduction, but not absence of crosslinked species and, importantly, monoglycyl-substituted oligomers were never found^17^.

A second study on the essentiality of *femAB* was done by Strandén and colleagues, who constructed a *femAB* mutant, AS145, by allelic replacement of the *femAB* operon by a tetracycline resistance marker^18^. AS145 showed impaired growth, methicillin hypersusceptibility, accumulation of monoglycyl substituted PG monomers and drastically reduced crosslinking of glycan strands, when compared to the parental strain^18^. *Cis*-complementation of the *femAB* mutation in AS145 with wild-type *femAB* restored synthesis of the pentaglycine crossbridge and methicillin resistance, but the growth rate remained low^19^. Therefore the authors postulated that survival of AS145 required compensatory or suppressor mutations^19^. Transcriptional analysis revealed that AS145 underwent severe metabolic adaptations to survive, including upregulation of membrane transporters associated with glycerol uptake (an osmoprotectant), upregulation of the arginine-deiminase pathway (an alternative for ATP production) and downregulation of nitrogen metabolism. Collectively these data suggested that *femAB* mutants adapted to survive with shortened crossbridges by drastically reducing metabolic activity to alleviate internal turgor^19^.

The *femAB* operon and the pentaglycine crossbridges are unique features of *S. aureus* among prokaryotes. This makes FemAB proteins potentially interesting targets for MRSA-specific drug design. In this work we wanted to investigate if full depletion of the *femAB* operon is lethal in an MRSA strain and to determine the phenotypic defects associated with lack of *femAB* expression.

## Results and Discussion

### The *femAB* operon is essential for the viability of *S. aureus*

Previous *S. aureus femAB* null mutants likely had compensatory mutations^16,17,19^. To evaluate the essentiality of *femAB* as well as the phenotypes resulting from FemAB depletion in a background without the existence of compensatory mutations, we constructed a conditional *femAB* mutant. The *femAB* operon of the clinically relevant CA-MRSA strain MW2 was placed under the control of the IPTG inducible P*spac* promoter. As the P*spac* promoter is known to be leaky^20^, a plasmid-encoded *lacI* repressor gene was provided to decrease the basal transcription of *femAB*. The resulting strain was named MW2-iFemAB.

Growth of MW2-iFemAB in liquid medium supplemented with IPTG at 0, 10, 25 and 500 µM was followed for 10 hours. In the presence of 500 µM of IPTG, growth of the conditional mutant was similar to the parental strain MW2 (Fig. 1a), therefore this concentration of inducer was used in subsequent assays. The growth rate of MW2-iFemAB decreased with decreasing IPTG concentrations and, surprisingly, no bacterial growth was observed in the absence of IPTG, indicating that this operon is essential for survival (Fig. 1a), contrarily to what was previously thought. We tested the essentiality of the *femAB* operon in an HA-MRSA background – strain COL – and likewise no growth of COL-iFemAB was observed in the absence of IPTG (Supplementary Fig. 1). To assess the effect of loss of FemAB activity on PG composition, we analysed the cell wall of MW2-iFemAB cells incubated with 500, 25 or 0 µM of IPTG until bacterial growth was arrested in the non-induced culture (see Methods). As expected, the muropeptide profiles in cells depleted of FemAB show a massive accumulation of peak 4 (see Supplementary Fig. 2 for peak assignment), which was previously identified as the monomeric pentapeptide substituted with a single glycine residue^21^, the substrate of FemA (Fig.1b, [IPTG] 0 µM). This was in contrast to cells where *femAB* expression was fully induced (Fig.1b, [IPTG] 500 µM), where the major monomeric form present was the pentaglycine substituted monomer (peak 5). Accordingly, lack of FemAB also prevented the formation of pentaglycine crosslinked forms such as dimers (peak 11), trimers (peak 15), tetramers (peak 16) and higher order forms, which co-elute near the end of the run (Fig. 1b, arrow). Low FemAB expression levels, just enough to sustain bacterial growth ([IPTG] 25 µM), resulted in the presence of some pentaglycine crosslinked forms (peaks 11, 15, 16, etc.). This degree of peptidoglycan structural organisation might be the minimum to ensure cell viability.

**Figure 1.**
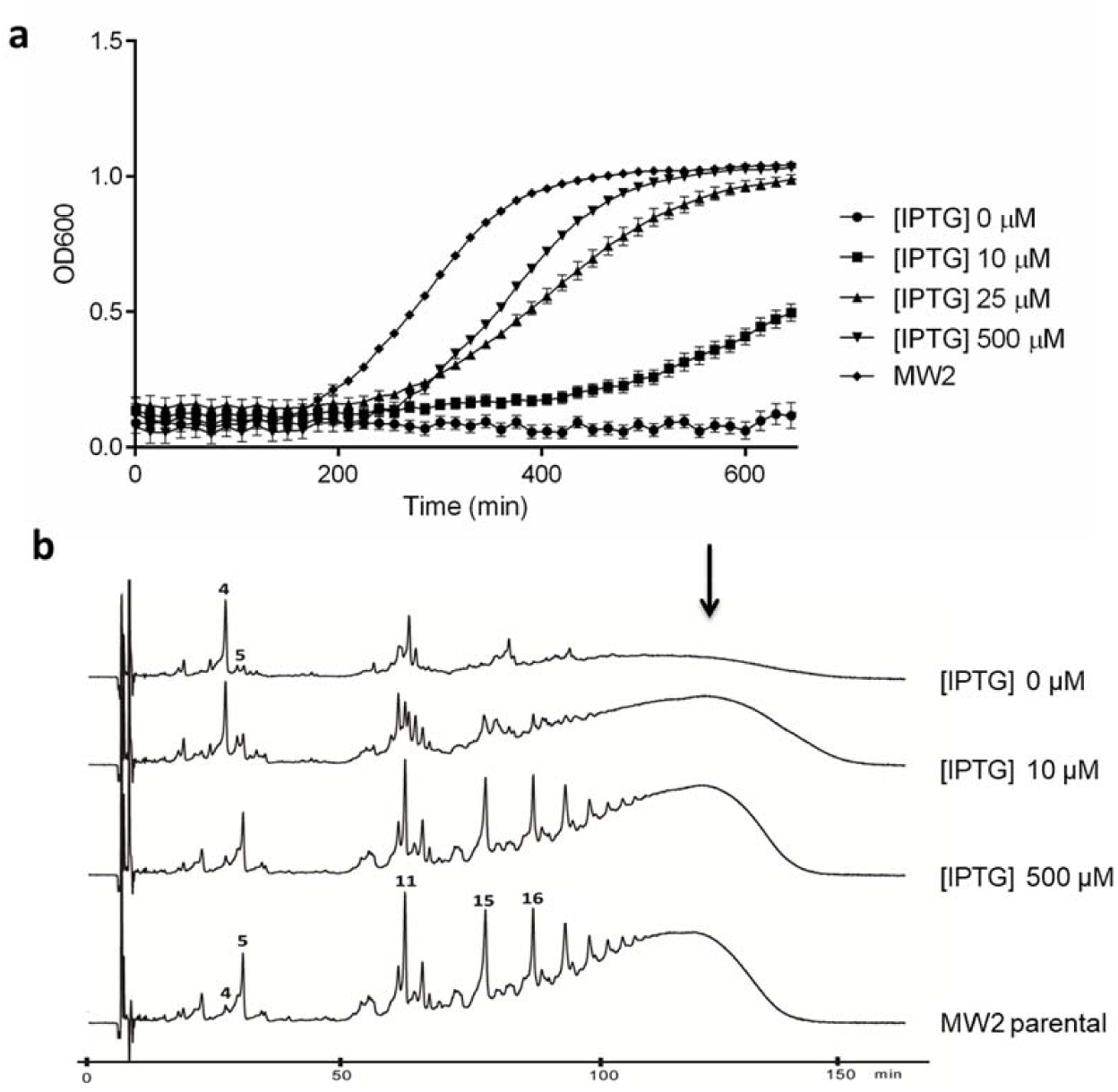
FemAB are essential for cell viability in *S. aureus*. **(a),** Growth curves of MW2-iFemAB strain with IPTG-inducible *femAB* operon. In the presence of high IPTG concentrations ([IPTG] 500 µM), growth was similar to the parental strain (MW2). Cell growth was reduced with decreasing IPTG concentrations. In the absence of IPTG ([IPTG] 0 µm), no cell growth was detected. Symbols indicate means and error bars indicate standard deviation from three biological replicates. **(b),** Muropeptide HPLC profiles of MW2-iFemAB grown in the presence of different levels of IPTG. Depletion of FemAB led to the accumulation of monomeric pentapeptides substituted with one glycine (peak 4), in contrast to pentaglycine forms (peak 5) seen in fully induced ([IPTG] 500 µM) or parental strain (MW2 parental) profiles (see Supplementary Fig. 2 for peak assignment). Loss of FemAB activity also impaired the formation of pentaglycine crosslinked forms such as di-, tri-and tetramers (peaks 11, 15 and 16, respectively) and higher order oligomers (black arrow). Muropeptide profiles shown are representative of three independent experiments.

### Loss of FemAB activity leads to membrane damage

The morphology of cells depleted of FemAB was analysed by super resolution structured illumination microscopy (SIM). MW2-iFemAB was grown with or without IPTG (500 µM). Following growth arrest of the non-induced culture (Supplementary Fig. 3, arrow), cells were stained with membrane dye FM 4-64, PG dye Van-FL and with DNA dye Hoechst 33342. In the presence of IPTG, MW2-iFemAB cells divided normally with DNA segregation preceding the synthesis of a division septum at mid-cell (Fig. 2a, top row). Cells containing multiple septa were rarely observed (<1%, N = 466 septating cells - Fig. 2b, left panel). In contrast, FemAB depleted cells often appeared as pseudomulticellular forms with two or more perpendicular septa (56%, N = 480 septating cells), suggesting that a second round of division starts before daughter cell separation is completed (Fig. 2b, right panel arrows). Furthermore, the nucleoid morphology of FemAB depleted cells was altered, with the presence of cells containing condensed DNA (Fig. 2a, bottom row asterisks). Our results are in agreement with previous reports that suppressed *fem* mutants show irregular placement of cross walls and retarded cell separation^16^. This phenotype can be a consequence of either multiple septation or defective splitting.

**Figure 2.**
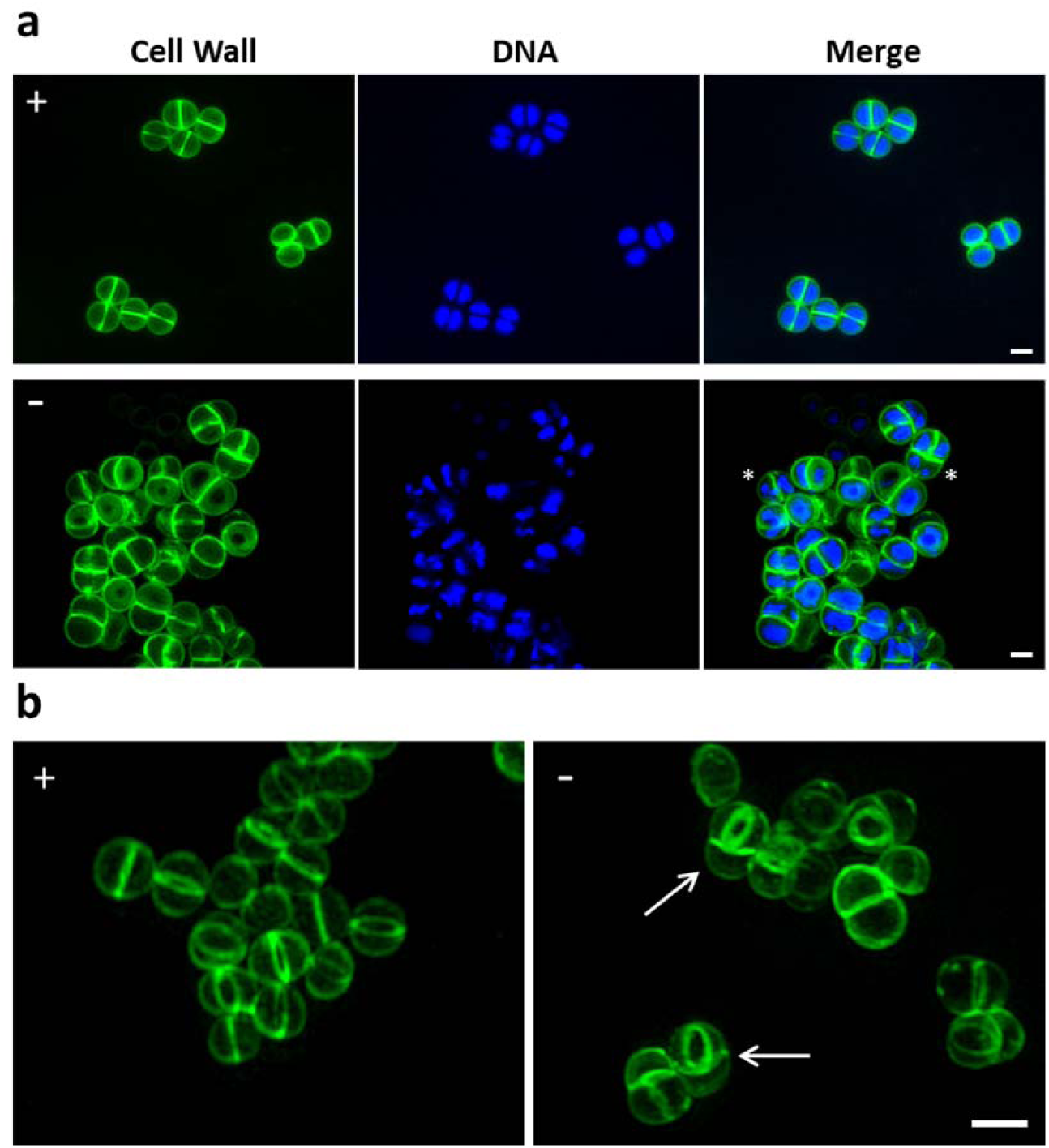
Loss of FemAB activity inhibits daughter cell separation during division. **(a)**, SIM images of MW2-iFemAB cells growing in the presence (+) or absence (-) of IPTG and labelled with cell wall dye Van-FL and DNA dye Hoechst 33342. **(b)**, 3D-SIM projections of MW2-iFemAB cells growing in the presence (+) or absence (-) of IPTG and stained with cell wall dye Van-FL. IPTG-induced cells divide normally with DNA segregation preceding the synthesis of the division septum at mid-cell (Panel **a**, top row and Panel **b**, left column). In contrast, FemAB depleted cells often had condensed nucleoids (Panel **a**, bottom row asterisks) and appeared as pseudo multicellular forms with two perpendicular septa (Panel **b**, white arrows). Scale bars, 1 µm

When cells depleted of FemAB were incubated for longer periods, we noticed a decrease in culture density, suggesting cell lysis (Supplementary Fig. 3). We therefore imaged cells 2 hours after growth arrest and observed extensive membrane damage, characterised by bulges and invaginations (Fig. 3a, arrow) and the presence of anucleate cells, indicative of loss of viability (Fig. 3a, asterisks). These results suggest that the inability of *S. aureus* to survive with shortened crossbridges could be because the three-dimensional structure of this alternate PG does not confer sufficient robustness and/or flexibility to bear the internal osmotic pressure, in these conditions, causing the cells to rupture. In order to test this hypothesis we incubated MW2-iFemAB in the absence of IPTG (to deplete FemAB expression) with increasing concentrations of NaCl added to the medium, to alleviate turgor. MW2-iFemAB was able to grow in the presence of NaCl in a dose dependent manner (Fig. 3b), confirming that in the absence of pentaglycine crosslinks, the PG layer is not able to withstand the internal pressure exerted on the membrane. This is in accordance with data from Hübscher and colleagues^19^, who showed by transcriptome analysis that *femAB* null mutant AS145 adapted to the FemAB deficit by tuning its metabolic pathways, presumably to reduce turgor. It is likely that monoglycine crossbridges are not suitable substrates for transpeptidation by *S. aureus* PBPs *in vivo* and thus crosslinking of glycan chains was stalled after FemAB depletion. Accordingly, solid-state NMR data obtained by Kim et al.^22^ indicated that monoglycyl crossbridges would be too short to connect the glycan chains of the *S. aureus* PG, and that crosslinking with such a reduced bridge length would require a major rearrangement of the tertiary structure of PG^22^.

**Figure 3.**
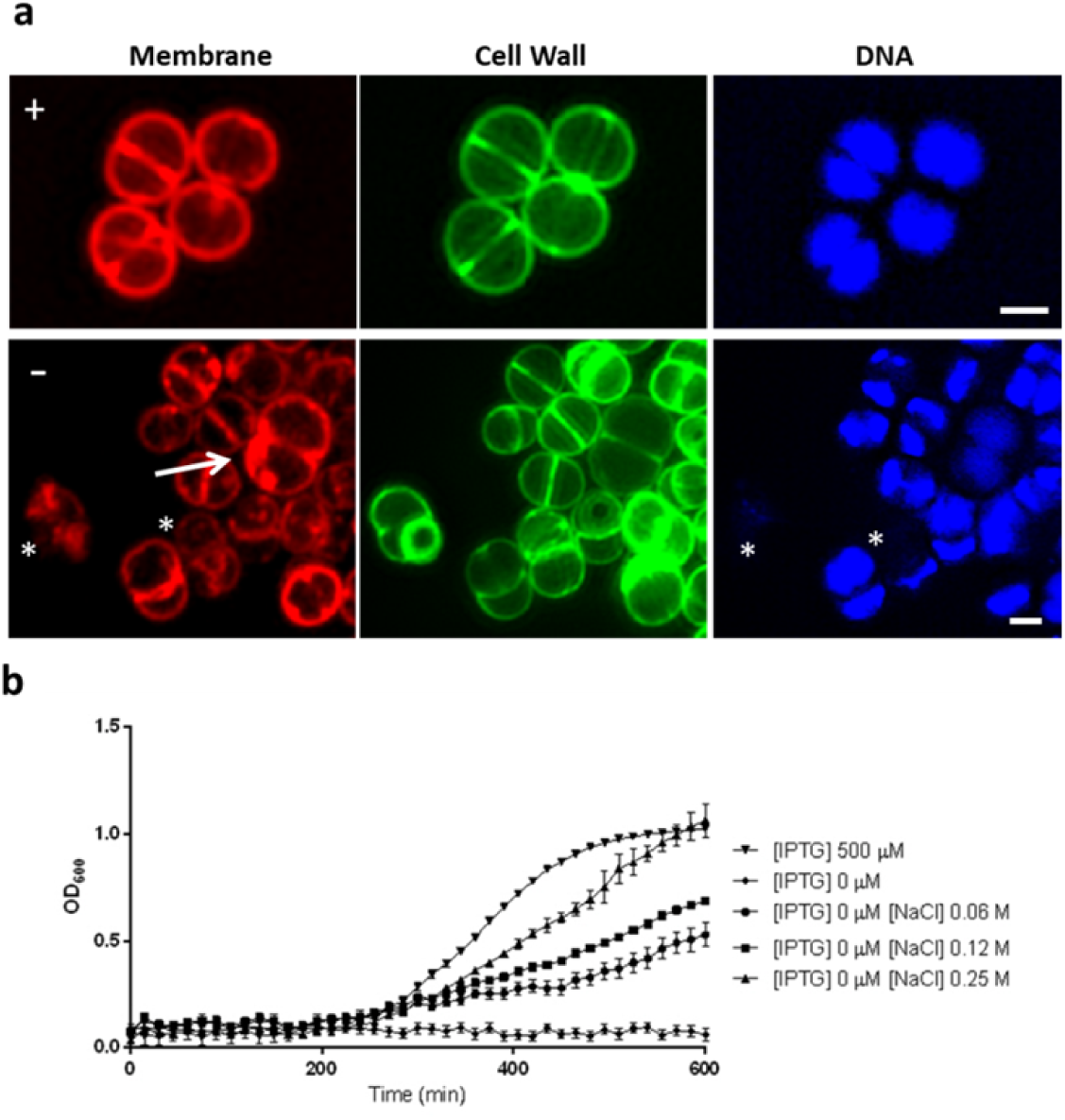
FemAB activity is required to withstand internal turgor. MW2-iFemAB cells depleted of FemAB for a period of 2 hours following growth arrest were stained with membrane dye FM 4-64, cell wall dye Van-FL and DNA dye Hoechst 33342 and imaged by SIM. FemAB depleted cells (**-**) show loss of membrane integrity characterised by bulging and invagination (white arrow) as well as absence of DNA staining (asterisks), indicative of loss of viability, when compared to IPTG-induced cells (**+**). Scale bars, 1 µm. (**b**), Growth rates of MW2-iFemAB incubated in the presence ([IPTG] 500 µM) or absence ([IPTG] 0 µM) of IPTG, or in the absence of IPTG with increasing NaCl concentrations. Addition of NaCl to the medium allowed cells to grow in the absence of FemAB expression, in a dose dependent manner. Symbols indicate means and error bars indicate standard deviation from three biological replicates.

### FemA activity is not required for membrane localisation

The Fem proteins are non-ribosomal peptidyl transferases which use dedicated amino acid charged tRNA molecules as substrates, an interesting activity seldom seen in nature^12^. The mechanism of this transfer is still poorly understood, as binding sites for entering tRNA molecules have not been identified. In the case of Fem proteins which transfer two amino acids, such as FemA (and FemB), the transfer of both glycines to lipid II appears to occur simultaneously rather than sequentially, judging from *in vitro* data, which may indicate that these proteins act as homodimers^23^. We have recently reported that all three Fem proteins of *S. aureus* localise to the membrane throughout the entire cell cycle, which was unexpected given that these proteins lack canonical transmembrane domains^24^. Therefore a possible mechanism of Fem localisation to the membrane could be through protein activity, which is dependent of interactions with both the substrate lipid II and glycyl-charged tRNA molecules.

In order to investigate the mechanism of localisation of FemA, we identified possible key regions in FemA required for activity, based on the known crystal structure of FemA^23^ and on homology to the FemX protein from *Weisella viridescens*^25,26^. We decided to focus on the putative transferase pocket that contains Arg220, Phe224 and Tyr327, which are conserved across the Fem family^23,26^. We also mined the sequence of FemA for regions which could bind DNA/RNA using DP-Bind^27,28^, in order to identify the putative tRNA-binding site. We found that the region with the highest probability of binding to RNA corresponded to the α6 helix (aa 176-188) of FemA, rich in Lys/Arg residues with polar and charged sidechains exposed to the solvent^23^, which could stabilise the entering tRNA. Specifically, amino acids Lys180 and Arg181 showed >96% probability of binding DNA/RNA in each of three individual prediction algorithms performed by DP-Bind (see Methods), and therefore were selected for mutagenesis.

To assess if the selected mutations had an effect on FemA transferase activity, we cloned wild-type *femA* into the pET-24b expression vector and performed site-directed mutagenesis on *femA* residues to obtain FemA mutants where the target residues were replaced by alanines. In this way, we constructed pET-FemA^RF220AA^ and pET-FemA^Y327A^, in order to express mutants in the transferase domain and pET-FemA^KR180AA^ to express a mutant of the predicted tRNA binding helix. We purified recombinant FemA^wt^, FemA^KR180AA^, FemA^RF220AA^ and FemA^Y327A^ with C-terminal histidine tags and synthesised the FemA substrate lipid II-Gly_1_ *in vitro* (see Methods). As recombinant FemA^RF220AA^ was very unstable and readily precipitated, we could not use it for further assays. Lipid II-Gly_1_ was trapped in Triton X-100 micelles and incubated with either FemA^wt^, FemA^KR180AA^ or FemA^Y327A^ in the presence of [U-^14^C]-glycine charged tRNA. After 30, 60 or 90 minutes the lipid fraction was extracted and separated by thin layer chromatography and radioactive glycine transfer to lipid II-Gly_1_ was measured. Both mutants FemA^KR180AA^ and FemA^Y327A^ transferred less [U-^14^C]-glycine to their substrate than FemA^wt^, consistent with a reduction of enzyme activity (Fig. 4a).

**Figure 4.**
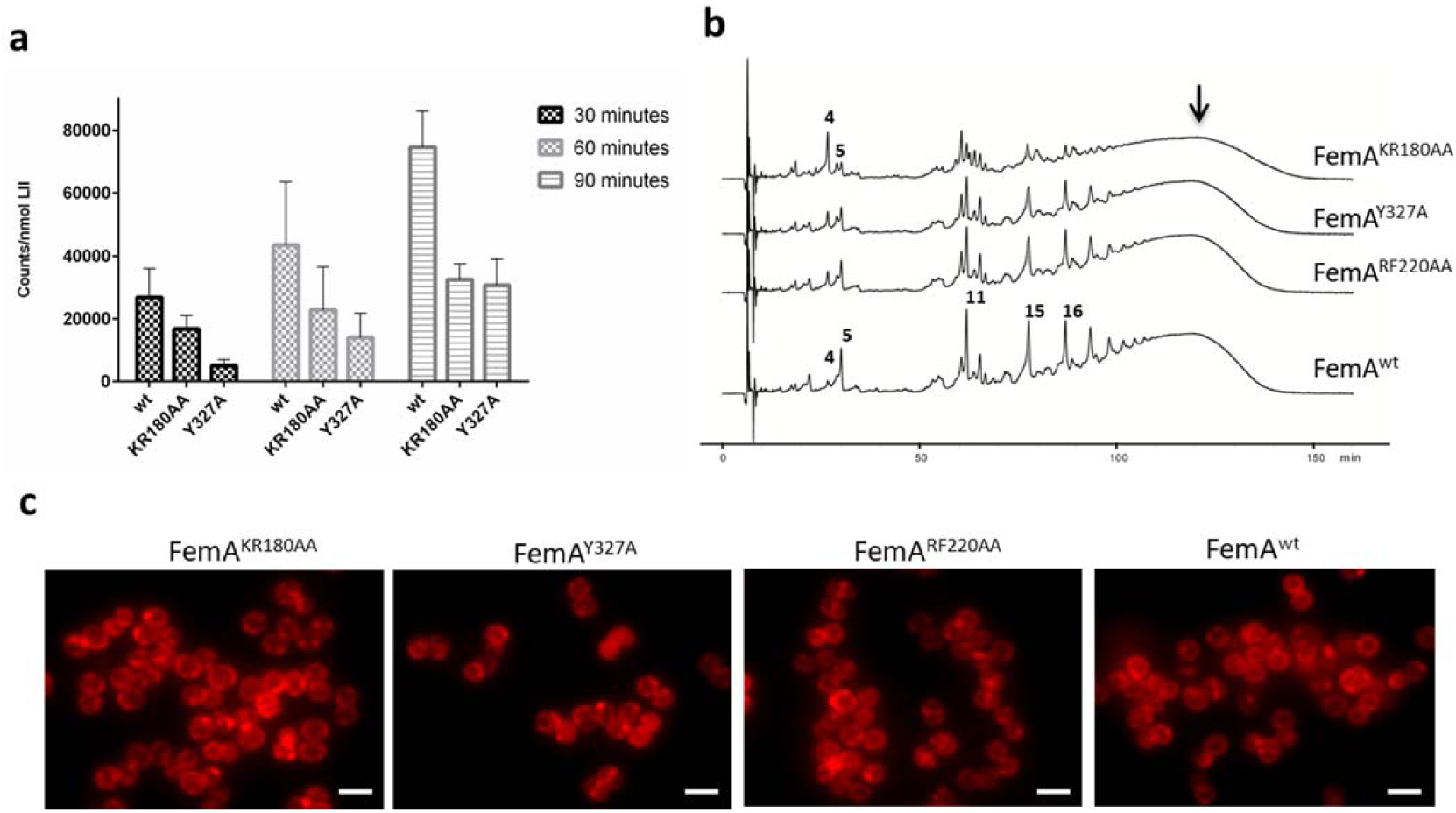
Selected mutations decrease FemA activity. **(a)**, Recombinant FemA^wt^, FemA^KR180AA^ and FemA^Y327A^ were incubated with lipid II-Gly_1_ in the presence of [U-^14^C]-glycine charged tRNA, for either 30, 60 or 90 minutes. Both FemA^KR180AA^ and FemA^Y327A^ showed decreased [U-^14^C]-glycine transfer to lipid II-Gly_1_ when compared to FemA^wt^, indicating reduced FemA activity. Columns denote mean values and error bars represent standard deviation from 3 independent experiments. **(b),** muropeptide profiles of MW2-iFemAB cells depleted of native FemAB expression and containing ectopically expressed wild-type FemA-mCherry and FemB (FemA^wt^) or derivatives with mutations in FemA-mCherry (FemA^KR180AA^, FemA^RF220AA^ and FemA^Y327A^) and wild-type FemB, from the cadmium-inducible promoter *Pcad*. Ectopic expression of FemA^wt^ complemented the lack of native FemAB expression, while expression of FemA^KR180AA^ led to accumulation of monoglycine monomer species (peak 4) with concomitant reduction in higher-order pentaglycine crosslinked species (peaks 11, 15, 16 and black arrow, see Supplementary Fig. 2 for peak assignment). Expression of FemA^Y327A^ or FemA^RF220AA^ led to similar phenotypes, albeit to a lesser extent. **(c)**, fluorescence microscopy images of strains described in (b). FemA^KR180AA^ localised to the membrane in >95% of the cells, similarly to FemA^wt^. FemA^Y327A^ and FemA^RF220AA^ appeared dispersed in the cytoplasm in a fraction of the population (27% and 10%, respectively, white arrows). N = 400 cells for each strain.

In order to both confirm loss of FemA activity *in vivo* and study protein localisation, we used the backbone of pFemAB^wt^, a replicative vector encoding a *femA-mCherry* fusion followed by *femB* (both under the control of a cadmium inducible promoter), to generate *femA-mCherry* alleles with the mutations described above. These expression plasmids were transformed into MW2-iFemAB, allowing us to deplete native *femAB* expression (in the absence of IPTG) and express mutant alleles (in the presence of cadmium).

We were able to complement the lack of *femAB* expression from the native locus by expressing *femA-mCherry-femB* from pFemAB^wt^ in the presence of cadmium (0.1 µM), as assessed by growth rates, morphology, lysostaphin (an enzyme that cuts pentaglycine bridges^29^) and oxacillin MICs and muropeptide composition (Table 1). Expression of the catalytic site mutants FemA^RF220AA^ and FemA^Y327A^ caused a reduction of the pentaglycine substituted monomer content in peptidoglycan (Fig. 4b), although morphology was similar to wild-type and no significant differences in lysostaphin and oxacillin MICs were observed (Table 1). In contrast, the double mutation in the α6 helix of FemA caused severe loss of FemA activity. The FemA^KR180AA^ mutant showed a marked reduction in growth rate, increased lysostaphin and decreased oxacillin resistances and a pseudomulticellular morphology when observed by microscopy (Table 1), similar to what was observed when depleting *femAB* expression. Furthermore, analysis of the muropeptide content in this mutant revealed a pronounced accumulation of monoglycyl substituted pentapeptides and concomitant reduction in pentaglycine crosslinked species (Fig. 4b and Table 1). Nevertheless, FemA^KR180AA^ still localised to the membrane, similarly to FemA^wt^, indicating that FemA localisation is independent of protein activity (Fig. 4c). Because loss of activity in FemA^KR180AA^ is likely due to a deficit in tRNA binding, it is possible that FemA^KR180AA^ could still localise to the membrane through recognition of the lipid-linked peptidoglycan precursor.

**Table 1.** *In vivo* activity profiles of FemA mutants.

|  | Doubling time<br>(min) | MIC lysostaphin<br>(µg/ml) | MIC oxacillin<br>(µg/ml) | Morphology | Gly5/Gly1<br>monomer<br>species fraction |
| --- | --- | --- | --- | --- | --- |
| MW2pFemA<sup>KR180AA</sup> | 52 | 2.5 | 0.4 | defective | 1:3 |
| MW2pFemA<sup>RF220AA</sup> | 30 | 0.15 | 1.6 | wt | 2:1 |
| MW2pFemA<sup>Y327A</sup> | 27 | 0.08 | 0.8 | wt | 2:1 |
| MW2pFemA<sup>wt</sup> | 27 | 0.08 | 3.2 | wt | 6:1 |
| Parental MW2 | 25 | 0.08 | 1.6 | wt | 6:1 |

The mutations in the substrate binding pocket had a minor effect on FemA localisation, since FemA^Y327A^ and FemA^RF220AA^ appeared dispersed in the cytoplasm in a small percentage of the cell population (Fig. 4c, white arrows). Nevertheless, as these mutations did not seem to decrease protein activity *in vivo* to a great extent, we could not conclude that the mechanism of FemA localisation to the membrane is via substrate recognition. An alternative possibility is that the recruitment of FemA to the membrane is mediated by protein-protein interactions with the membrane associated eukaryotic-type serine/threonine kinase Stk, a global cell wall synthesis regulator, which was recently shown to interact with FemA and FemB by bacterial two hybrid^30^. However, the same study could not find interactions between Stk and FemX, which initiates Lipid II crossbridge synthesis, or between FemX and FemA/B^30^. Further experiments are necessary to clarify the interactions of Fem proteins with each other and with their substrates, in order to understand how the localisation and timing of PG crossbridge synthesis is modulated during the cell cycle.

### Concluding remarks

The structural features of the staphylococcal PG seem remarkably unique in nature, as pentaglycine crosslinks have not been observed outside of the genus. These long bridges likely confer high flexibility to *S. aureus* PG that allows a high level of PG crosslinking, which in turn allows the cell to withstand high internal turgor. Accordingly, *femAB* mutants isolated in the past adapted to life with shortened crossbridges by drastically reducing metabolic activity^18^. Moreover, the nature and length of PG branching has been implicated in resistance to β-lactams, not only in *S. aureus* but also in other bacteria such as *Streptococcus pneumoniae*^7,31-33^.

We have shown that the depletion of the *femAB* operon is lethal in CA-MRSA strain MW2 and in HA-MRSA strain COL, leading to the disruption of the cell envelope, causing cells to lose viability. This suggests that monoglycyl-substituted muropeptides are not good substrates for transpeptidation *in vivo*, either because transpeptidases fail to recognise them or because different *S. aureus* glycan strands are too far apart to be crosslinked via crossbridges with only one glycine.

## Methods

### Bacterial growth conditions

Strains and plasmids constructed for this study are listed in Supplementary Table 1. *S. aureus* strains were grown in tryptic soy broth (TSB, Difco) at 200 r.p.m with aeration at 37 °C or on tryptic soy agar (TSA, Difco) at 30 or 37 °C. *Escherichia coli* strains were grown in Luria– Bertani broth (Difco) with aeration, or Luria–Bertani agar (Difco) at 37 or 30 °C. When necessary, antibiotics ampicillin (100 μg/ml), erythromycin (10 μg/ml), kanamycin (50 μg/ml), neomycin (50 μg/ml) or chloramphenicol (30 μg/ml) were added to the media. Unless stated otherwise, isopropyl β-D-1-thiogalactopyranoside (IPTG, Apollo Scientific) was used at 500 µM to induce expression of constructs under the control of the P*spac* promoter. Cadmium chloride (Sigma-Aldrich) was used at 0.1 µM when required to induce expression of constructs under the control of the P*cad* promoter.

### Construction of *S. aureus* strains

In order to construct an *S. aureus* strain with the *femAB* operon under the control of an inducible promoter, a fragment containing the first 400 bp of *femA* was amplified from *S. aureus* MW2 DNA with primer pair spacfemab_P1 EcoRI/spacfemab_P2 BamHI (see Supplementary Table 2 for primers sequences), cut with EcoRI and BamHI restriction enzymes and cloned into pMUTIN4^34^, downstream of the P*spac* promoter, giving plasmid pFemABi, which was sequenced. pFemABi was then propagated in DC10B cells, electroporated into electrocompetent RN4220 cells, and transduced (using phage 80α) to MW2 and COL, where it integrated in the *femAB* locus by homologous recombination. The resulting strain contains a truncated copy of *femA* under the control of the *femAB* native promoter, and the *femAB* operon under the control of P*spac*. Multicopy plasmid pMGPII^35^, which encodes P*spac* repressor LacI, was then transduced into this strain, giving rise to MW2-iFemAB and COL-iFemAB.

To construct *S. aureus* strains expressing mutated alleles of FemA-mCherry together with wild-type FemB, first a *femA-mCherry-STOP* codon*-femB* was amplified from pMADfemAmch^24^ using primers pcnfemab_P1 BamHI and pcnfemab_P2 EcoRI. This fragment was cut with BamHI and EcoRI and cloned into replicative vector pCNX^36^, under the control of P*cad*, giving plasmid pFemAB^wt^. pFemAB^wt^ was then used as the template for site-directed mutagenesis using Phusion polymerase (Thermo Scientific) following manufacturer’s instructions. Primers fema_kr180aa_fw/ fema_kr180aa_rev were used to generate pFemA^KR180AA^, encoding both K180A and R181A mutations; Primers fema_rf220aa_fw/ fema_rf220aa_rev were used to generate pFemA^RF220AA^, encoding both R220A and F224A mutations; and primers fema_Y327a_fw/ fema_Y327a_rev were used to generate pFemA^Y327A^, encoding the Y327A mutation. Each plasmid was sequenced to confirm the presence of the mutations. pFemAB^wt^, pFemA^KR180AA^, pFemA^RF220AA^ and pFemA^Y327A^ were propagated in DC10B, electroporated into RN4220 and transduced to MW2-iFemAB, giving strains MW2pFemAB^wt^, MW2pFemA^KR180AA^, MW2pFemA^RF220AA^ and MW2pFemA^Y327A^, respectively.

### Growth curves of *S. aureus* strains

To assess whether the *femAB* operon is essential for viability, or the effects of FemA mutations on growth rates, overnight cultures of MW2, MW2-iFemAB, COL, COL-iFemAB, MW2pFemAB^wt^, MW2pFemA^KR180AA^, MW2pFemA^RF220AA^ and MW2pFemA^Y327A^ grown in TSB with 500 µM of IPTG, with the appropriate antibiotics (when applicable, at the concentrations described above) were back-diluted 1:500 in the same medium and grown until the cultures reached an OD_600_ of 0.7. At this point the cultures were washed three times to remove IPTG and back-diluted to an OD_600_ of 0.007 in fresh TSB containing either 0, 10, 25 or 500 µM of IPTG or 0.06, 0.12 or 0.25 of NaCl, in the case of MW2-iFemAB and COL-iFemAB. In the case of MW2pFemAB^wt^, MW2pFemA^KR180AA^, MW2pFemA^RF220AA^ and MW2pFemA^Y327A^, cells were incubated without IPTG and cadmium chloride was added at 0.1 µM to drive the expression of either wild-type or mutant *femA* alleles from the pCNX-based plasmids.

Growth of all strains was monitored for 10 hours in a Bioscreen C Analyzer (Growth Curves USA), at 37 °C with shaking with OD_600_ readings taken every 15 minutes. Growth curves were obtained from three independent experiments done with three biological replicates.

### Minimum inhibitory concentration (MIC) assays

MICs of lysostaphin and oxacillin were determined by broth microdilution in sterile 96-well plates. The medium used was TSB, containing a series of two-fold dilutions of each compound. Cultures of *S. aureus* strains and mutants were added at a final density of ∼5×10^5^ CFU ml^−1^ to each well. Wells were reserved in each plate for sterility control (no cells added) and cell viability (no compound added). Plates were incubated at 37°C. Endpoints were assessed visually after 48 h and the MIC was determined as the lowest concentration that inhibited growth. All assays were done in triplicate.

### Purification and analysis of *S. aureus* muropeptides

To evaluate changes in the peptidoglycan composition caused by the depletion of the *femAB* operon, or caused by the expression of mutant FemA proteins, cells of MW2, MW2-iFemAB, MW2pFemA^wt^, MW2pFemA^KR180AA^, MW2pFemA^RF220AA^ and MW2pFemA^Y327A^ were first grown overnight in TSB supplemented with 500 µM of IPTG and the applicable antibiotics. Cultures were then washed three times to remove the IPTG and back-diluted 1:500 in fresh TSB with 0, 10 or 500 µM of IPTG, in the case of MW2 and MW2-iFemAB, or in the presence of cadmium chloride and absence of IPTG, in the case of MW2pFemA^wt^, MW2pFemA^KR180AA^, MW2pFemA^RF220AA^ and MW2pFemA^Y327A^. Cells were collected at mid-exponential phase and PG was purified as described by Filipe et al.^37^. Muropeptides were prepared from PG samples by digestion with mutanolysin (0.135 U/µg of PG, from Sigma-Aldrich) and analysed by reverse phase HPLC using a Hypersil ODS (C18) column (Thermo-Fisher Scientific). Muropeptide species were eluted in 0.1 M sodium phosphate, pH 2.0, with a gradient of 5–30% methanol for 155 minutes and detected at 206 nm.

### *S. aureus* imaging by fluorescence microscopy

To evaluate changes in morphology caused by the depletion of the *femAB* operon, or to investigate the localisation of mutant FemA proteins, cells of MW2, MW2-iFemAB, MW2pFemA^wt^, MW2pFemA^KR180AA^, MW2pFemA^RF220AA^ and MW2pFemA^Y327A^ were first grown overnight in TSB supplemented with 500 µM of IPTG and the applicable antibiotics. Cultures were then washed three times to remove the IPTG and back-diluted 1:500 in fresh TSB with 0, 25 or 500 µM of IPTG, in the case of MW2 and MW2-iFemAB, or in the presence of cadmium chloride and absence of IPTG, in the case of MW2pFemA^wt^, MW2pFemA^KR180AA^, MW2pFemA^RF220AA^ and MW2pFemA^Y327A^. Cells were grown to an OD600 nm of 0.4-0.6, harvested and then washed with phosphate buffer saline (PBS). Subsequently, cells were washed with PBS, mounted on microscope slides covered with a thin layer of 1% agarose in PBS and imaged by fluorescence microscopy.

To evaluate defects in cell morphology, cells were incubated with membrane dye Nile Red (10 µg/ml, Invitrogen), Hoechst 33342 (10 µg/ml, Invitrogen) and a mixture containing equal amounts of vancomycin (Sigma) and a BODIPY FL conjugate of vancomycin (Van-FL, Molecular Probes) to a final concentration of 0.8 µg/ml, for 5 minutes at room temperature. Cells were then washed three times with PBS before being spotted on the agarose pads. super-resolution Structured Illumination Microscopy (SIM) imaging was performed using an Elyra PS.1 microscope (Zeiss) with a Plan-Apochromat 63x/1.4 oil DIC M27 objective. SIM images were acquired using five grid rotations, unless stated otherwise, with 34 μm grating period for the 561 nm laser (100 mW), 28 μm period for 488 nm laser (100 mW) and 23 μm period for 405 nm laser (50 mW). Images were captured using a Pco.edge 5.5 camera and reconstructed using ZEN software (black edition, 2012, version 8.1.0.484) based on a structured illumination algorithm, using synthetic, channel specific optical transfer functions and noise filter settings ranging from −6 to −8.

Wide-field fluorescence microscopy was performed using a Zeiss Axio Observer microscope with a Plan-Apochromat 100×/1.4 oil Ph3 objective. Images were acquired with a Retiga R1 CCD camera (QImaging) using Metamorph 7.5 software (Molecular Devices).

### Overexpression and purification of recombinant His-tagged proteins

Recombinant proteins were purified essentially as described by Rohrer et al.^38^, with some modifications. Single colonies of BL21 (DE3) expression strains containing either plasmid pFemAB^wt^, pFemA^KR180AA^B, pFemA^RF220AA^B or pFemA^Y327A^B were isolated from LA plates with kanamycin and used to inoculate LB (1 L) containing kanamycin. Cultures were grown to an OD_600nm_ of approximately 0.6 at which point IPTG was added (final concentration 1 mM) and incubated for 3 hours with shaking (150 rpm) at 30 °C. Cells were harvested by centrifugation and washed with 50 mM sodium phosphate buffer (pH 7.5) containing 300 mM NaCl and 20% glycerol. Afterwards, cells were suspended in the same buffer, containing PMSF (final concentration, 0.1 mM) and lysozyme (final concentration, 1 mg/mL), and incubated on ice for 30 min. Cells were then disrupted three times in an ultrasonicator and centrifuged for 30 min at 4°C to precipitate cell debris. The resulting supernatant was purified by affinity chromatography using a Ni-NTA column (Qiagen), following manufacturer’s instructions. Protein concentration was assessed using a BCA Protein Assay Kit (Pierce).

### Synthesis and purification of lipid II and lipid II-Gly_1_

Lipid II was prepared by reacting undecaprenyl phosphate (Larodan), UDP-MurNAc-pentapeptide from *Staphylococcus simulans*, UDP-GlcNAc (Sigma) and membrane proteins of *Micrococcus luteus* as previously described^14^. Monoglycyl lipid II was synthesised by reacting lipid I with tRNA preparations, in the presence of enzymes FemX and GlyS, according to the method described by Schneider et al.^14^. Lipid intermediates were extracted from reaction mixtures with an equal volume of butanol/pyridine acetate (2:1; vol:vol; pH 4.2). Extracts were then purified by anion-exchange chromatography using a Hi-Trap DEAE FF-agarose column (Amersham Biosciences) by reverse-phase HPLC and eluted in a linear gradient from chloroform–methanol–water (2:3:1) to chloroform–methanol-300 mM ammonium bicarbonate (2:3:1). The fractions containing lipid species were identified by thin layer chromatography with chloroform–methanol–water–ammonia (88:48:10:1) as solvent^39^. The concentration of purified lipids was calculated by measuring inorganic phosphates released after the treatment with perchloric acid, as described previously^40^.

### FemA enzymatic activity assay

In order to compare the activity of wild-type FemA to selected FemA mutants, enzymatic reactions were performed as described previously^14^. Briefly, 100 µl reactions were prepared containing 2.5 nmol of lipid II-Gly_1_, 10 µg of glycyl-tRNA synthetase (GlyS), 25 µg of tRNA, 2 mM ATP and 50 nmol [^14^C]-glycine in Tris buffer (100 mM Tris-HCl, 20 mM MgCl_2_, pH 7.5, and 0.8% Triton X-100). Then 2.7 µg of wild-type FemA or FemA mutant protein was added and the reaction mixtures were incubated for 30, 60 or 90 minutes at 30°C. Lipid intermediates were then extracted and analysed by thin layer chromatography, as described above. Finally, the amount of [^14^C]-glycine transferred to lipid II-Gly_1_ was quantified using phosphoimaging in a STORM system (GE Healthcare). Enzymatic assays were done in triplicate.

### Identification of FemA residues possibly involved in tRNA binding

Identification of FemA residues which could bind glycyl-charged tRNA was performed using DP-Bind^27,28^ (http://lcg.rit.albany.edu/dp-bind/), a sequence-based web server which predicts DNA/RNA binding domains in proteins based on biochemical properties of amino acids and evolutionary information. Probability maps were generated using PSI-BLAST position-specific scoring matrix (PSSM) and three distinct machine learning methods that use evolutionary information: support vector machine (PSSM-SVM), kernel logistic regression (PSSM-KLR), and penalized logistic regression (PSSM-PLR). FemA residues K180 and R181 were identified as possibly part of DNA-binding domains based on strict consensus between the three methods. SwissPdb viewer/Deep view (http://www.expasy.org/spdbv/) was used to evaluate the structure of FemA, using file 1LRZ (doi: 10.2210/pdb1LRZ/pdb) deposited in the RCSB PDB by Benson et. al.

## Supporting information

## Acknowledgments

This study was funded by the European Research Council through grant ERC-2017-COG 771709 (to M.G.P.), by Project LISBOA-01-0145-FEDER-007660 Microbiologia Molecular, Estrutural e Celular (to ITQB-NOVA), by the German Research Foundation (DFG; SCHN1284/1-2) to T.S. and FCT fellowship SFRH/BD/71993/2010 (J.M.M.).

## Author Contributions

J.M.M., H.G.S. and M.G.P. designed the research. J.M.M. constructed all strains and performed all experiments with the exception of the glycine incorporation assays, which were performed by D.M. J.M.M., D.M., T.S., S.R.F. and M.G.P. analysed the data. J.M.M. and M.G.P. wrote the manuscript.

## Data Availability

Source data are available from the corresponding author upon reasonable request.

