## Supplementary material for "The pentaglycine bridges of *Staphylococcus aureus* peptidoglycan are essential for cell integrity"

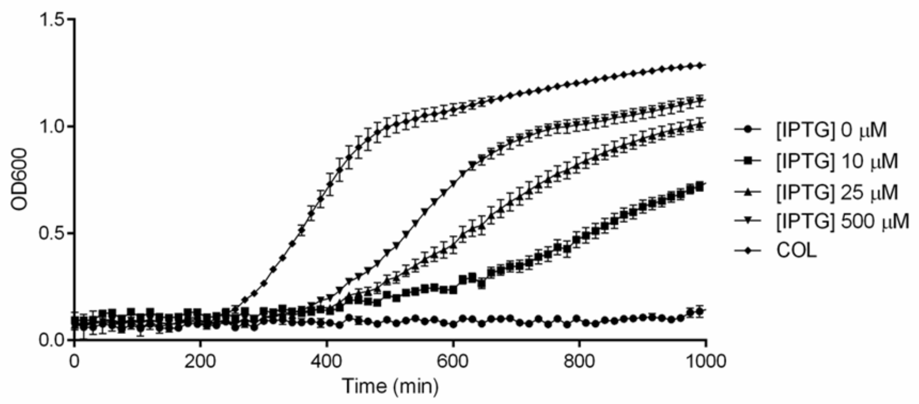


**Supplementary Figure 1. *femAB* is essential in COL.** Growth curves of COL-iFemAB with IPTG-inducible *femAB* operon. In the absence of IPTG ([IPTG] 0 μM), no growth was detected. Growth was rescued with increasing concentrations of IPTG. Symbols indicate means and error bars indicate standard deviation from three biological replicates.

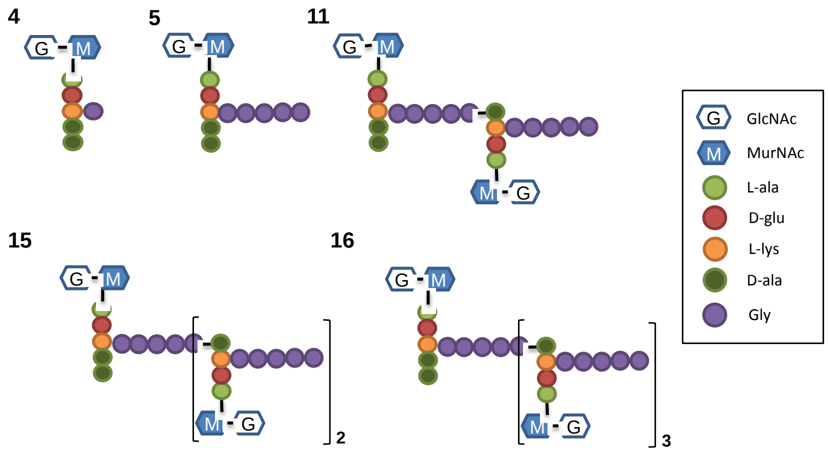


**Supplementary Figure 2. Chemical structures of muropeptide species.** Proposed structures of the peaks present in muropeptide chromatograms, according to de Jonge and Tomasz[1]. Muropeptides are numbered according to increasing retention times. GlcNAc – N-acetylglucosamine, MurNAc – N-acetylmuramic acid.

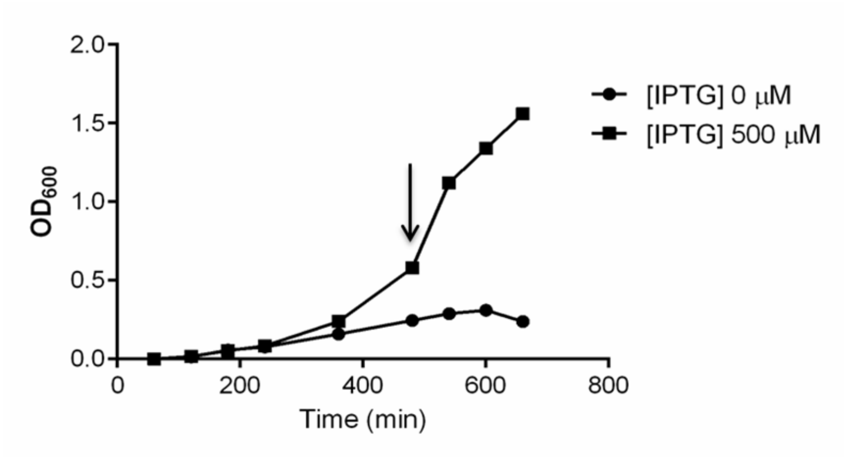


**Supplementary Figure 3. Depletion of FemAB led to growth arrest**. Growth curves of MW2-iFemAB in the presence ([IPTG] 500 µM) or absence ([IPTG] 0 µM) of IPTG, to determine the timing of growth arrest in the non-induced condition (see Methods). Cells were collected for microscopy at the indicated time point (black arrow).

**Supplementary Table 1 – Strains and plasmids used in this study.**

| Strains | Description | Source or reference |
|---|---|---|
| *E. coli* | | |
| DC10B | Δ*dcm* in the DH10B background; Dam methylation only | 2 |
| BL21 (DE3) | B $F^-$ *dcm ompT hsdS*($r_B^-$ $m_B^-$*) gal* λ(DE3) | Stratagene |
| BL21-FemX | BL21(DE3) expressing full length C-ter His-tagged FemX; $Kan^r$ | This work |
| BL21-GlyS | BL21(DE3) expressing full length C-ter His-tagged GlyS; $Kan^r$ | This work |
| BL21-$FemA^{wt}$ | BL21(DE3) expressing full length C-ter His-tagged FemA; $Kan^r$ | This work |
| BL21-$FemA^{KR180AA}$ | BL21(DE3) expressing mutant C-ter His-tagged $FemA^{KR180AA}$; $Kan^r$ | This work |
| BL21-$FemA^{RF220AA}$ | BL21(DE3) expressing mutant C-ter His-tagged $FemA^{RF220AA}$; $Kan^r$ | This work |

| Strains | Description | Source or reference |
|---|---|---|
| BL21-FemA$^{Y327A}$ | BL21(DE3) expressing mutant C-ter His-tagged FemA$^{Y327A}$; Kan$^{r}$ | This work |
| *S. aureus* | | |
| RN4220 | Restriction-deficient derivative of NCTC8325-4 | 3 |
| MW2 | CA-MRSA; SCCmec type IV, ST1, pbla$^{+}$ | 4 |
| MW2-iFemAB | MW2 *femAB::*pMUTIN*Pspac-femAB* pMGPII; Ery$^{r}$ Cm$^{r}$ | This work |
| MW2pFemA$^{wt}$ | MW2 *femAB::*pMUTIN*Pspac-femAB* pMGPII pFemAB$^{wt}$; Ery$^{r}$ Cm$^{r}$ Kan$^{r}$ | This work |
| MW2pFemA$^{KR180AA}$ | MW2 *femAB::*pMUTIN*Pspac-femAB* pMGPII pFemA$^{KR180AA}$B; Ery$^{r}$ Cm$^{r}$ Kan$^{r}$ | This work |
| MW2pFemA$^{RF220AA}$ | MW2 *femAB::*pMUTIN*Pspac-femAB* pMGPII pFemA$^{RF220AA}$B; Ery$^{r}$ Cm$^{r}$ Kan$^{r}$ | This work |
| MW2pFemA$^{Y327A}$ | MW2 *femAB::*pMUTIN*Pspac-femAB* pMGPII pFemA$^{Y327A}$B; Ery$^{r}$ Cm$^{r}$ Kan$^{r}$ | This work |
| COL | HA-MRSA | 5 |
| COL-iFemAB | COL *femAB::* pMUTIN*Pspac-femAB* pMGPII; Ery$^{r}$ Cm$^{r}$ | This work |

| Plasmids | Description | Source or reference |
|---|---|---|
| pMUTIN4 | *S. aureus* integrative vector containing an IPTG-inducible P*spac* promoter; Amp$^{r}$ Ery$^{r}$ | 6 |
| pMGPII | *S. aureus* replicative plasmid containing *lacI*; Amp$^{r}$ Cm$^{r}$ | 7 |
| pCNX | Replicative vector containing a cadmium inducible P*cad* promoter; Amp$^{r}$ Kan$^{r}$ | 8 |
| pFemABi | pMUTIN4 derivative containing a *femA* DNA fragment under the control of P*spac;* Amp$^{r}$ Ery$^{r}$ | This work |
| pMADfemAmch | Vector containing a *femA-mCherry-STOP-femB* DNA fragment; Amp$^{r}$ Ery$^{r}$ | 9 |
| pFemAB$^{wt}$ | pCNX derivative expressing a FemA-mCherry fusion and FemB, both under the control of P*cad;* Amp$^{r}$ Kan$^{r}$ | This work |
| pFemA$^{KR180AA}$B | pCNX derivative expressing a mutant FemA(K180A, R181A)-mCherry fusion and FemB, both under the control of P*cad;* Amp$^{r}$ Kan$^{r}$ | This work |
| pFemA$^{RF220AA}$B | pCNX derivative expressing a mutant FemA(R220A, F224A)-mCherry fusion and FemB, both under the control of P*cad;* Amp$^{r}$ Kan$^{r}$ | This work |
| pFemA$^{Y327A}$B | pCNX derivative expressing a mutant FemA(Y327A)-mCherry fusion and FemB, both under the control of P*cad;* Amp$^{r}$ Kan$^{r}$ | This work |
| pET-24b | *E. coli* replicative vector for the expression of proteins with a $His_6$ fusion at the C-terminus, under P*spac* promoter; Kan$^{r}$ | Novagen |

| Plasmids | Description | Source or reference |
|---|---|---|
| pET-GlyS | pET-24b derivative encoding a GlyS-$His_6$ fusion; $Kan^r$ | 10 |
| pET-FemX | pET-24b derivative encoding a FemX-$His_6$ fusion; $Kan^r$ | 10 |
| pET-$FemA^{wt}$ | pET-24b derivative encoding a FemA-$His_6$ fusion; $Kan^r$ | This work |
| pET-$FemA^{KR180AA}$ | pET-24b derivative encoding a mutant FemA(K180A, R181A)-$His_6$ fusion; $Kan^r$ | This work |
| pET-$FemA^{RF220AA}$ | pET-24b derivative encoding a mutant FemA(R220A, F224A)-$His_6$ fusion; $Kan^r$ | This work |
| pET-$FemA^{Y327A}$ | pET-24b derivative encoding a mutant FemA(Y327A)-$His_6$ fusion; $Kan^r$ | This work |

Amp, ampicillin; Kan, kanamycin; Ery, erythromycin; Cm, chloramphenicol

**Supplementary Table 2. Oligonucleotides used in this study.**

| Primer Name | Sequence (5'-3') |
|---|---|
| spacfemab_P1 EcoRI | GCGCGAATTCATGAAGTTTACAAATTTAACAGC |
| spacfemab_P2 BamHI | CGCGGGATCCTTATCAAAGAACCAATCATTACCAGCATTAC |
| pcnfemab_P1 BamHI | GCGCGCGGATCCGCAAATACGGAAATGAAATTAATTAACG |
| pcnfemab_P2 EcoRI | CGCGCGGAATTCCTATTTCTTTAATTTTTTACGTAATTTATC |
| fema_kr180aa_fw | ATGGACTTAGAGCAGCAAACACGAAAAAAGTTAAAAAGAATG |
| fema_kr180aa_rev | TTTTCGTGTTTGCTGCTCTAAGTCCATCCATATTTTTAATGATG |
| fema_rf220aa_fw | GCTGATGCTGATGACAAAGCTTACTACAATCGCTTAAAATATTAC |
| fema_rf220aa_rev | GTAGTAAGCTTTGTCATCAGCATCAGCAAAAGCTTTTGATTC |
| fema_Y327a_fw | GAAGTTGTTGCTTATGCTGGTGGTACATCAAATGCATTCC |
| fema_Y327a_rev | ACCAGCATAAGCAACAACTTCAAATGGATTGATAAAGAAG |
| femaexpress_P1 BamHI | CGCGCGGATCCATGAAGTTTACAAATTTAACAGCTAAAGAGTTTG |
| femaexpress_P2 EcoRI | CGCGCGAATTCCTAAAAAAATTCTGTCTTTAACTTTTTTAAGTGC |

Underlined sequences correspond to restriction sites
